# Opiate anticipation, opiate induced anatomical changes in hypocretin (Hcrt, orexin) neurons and opiate induced microglial activation are blocked by the dual Hcrt receptor antagonist suvorexant, while opiate analgesia is maintained

**DOI:** 10.1101/2023.09.22.559044

**Authors:** Ronald McGregor, Ming-Fung Wu, Thomas C. Thannickal, Jerome M. Siegel

## Abstract

We previously found that heroin addiction in humans is accompanied by an increase in the number of detected Hcrt neurons and a decrease in their soma size. We now show that the increased number of Hcrt cells visible after morphine treatment is likely the result of increased Hcrt production in neurons having sub-detection levels of the peptides. We find that morphine increases Hcrt projections to the ventral tegmental area (VTA), the level of tyrosine hydroxylase enzyme (TH) and the number of TH positive cells in VTA, with no changes in the adjacent substantia nigra. We find that the dual Hcrt receptor antagonist suvorexant prevents morphine-induced changes in the number and size of Hcrt neurons, microglial activation and morphine anticipatory behavior, but does not diminish morphine analgesia. These findings suggest that combined administration of opiates and suvorexant may be a less addictive way of administering opiates for pain relief in humans.

---

The annual US rate of opioid overdose deaths now exceeds 76,000, much greater than the annual rates for automobile or gun deaths (CDC website). This is in contrast to the annual US opioid overdose death rate of 8,000 recorded before 1990. Of those who began abusing opioids in the 2000s, 75 percent reported that their first opioid was prescribed for the relief of pain^1 2^. This progressed to illegal opioid pill acquisition or to heroin or fentanyl use^3^-^5^. Although non-opioid analgesics can be used for relatively minor pain, severe burns, cancer, joint inflammation, sickle cell disease, bone damage and many other painful conditions often cannot be effectively treated with non-opioid analgesics. These disorders cause immense suffering.

We^6^-^10^ and others^11^ have demonstrated that increased neuronal discharge in hypocretin (Hcrt, orexin) neurons is linked to the performance of rewarded tasks in wild type (WT) mice, rats, cats and dogs. Mice in which the Hcrt peptide is genetically knocked out (Hcrt-KOs) learn a bar press task for food or water as quickly as their WT littermates. However, when the effort to obtain the reward is increased in a “progressive ratio,” they all quit the task within 1 h, whereas all their WT littermates continue bar pressing until the end of the 2 h test period. In contrast, the Hcrt-KOs perform as well as WT controls on progressive ratio avoidance tasks, suggesting an emotional specificity in their response deficit^6^. Normal dogs playing in a yard have a large increase in cerebrospinal fluid Hcrt level. But when these same dogs are made to run on a treadmill, there is no change in Hcrt level, despite similar elevations of heart rate, respiratory rate and blood pressure^8^. We found that Hcrt is released in the brain of humans when they are engaged in tasks they enjoy, but not when they are aroused by pain or when they are feeling sad^12^.

Dopamine neurons, particularly those located in the ventral tegmental area (VTA) are known to play a significant role in reinforcement in general and in opiate use disorder (OUD) in particular^13-15^. Hcrt and dopamine are evolutionarily linked from both a neurochemical and anatomical perspective^16^. VTA plasticity associated with drug rewards requires functional Hcrt receptors^17^. The levels of dopamine and its major metabolites in the nucleus accumbens are markedly increased by the microinjection of Hcrt into the VTA^18, 19^. Hcrt neurons project strongly to the nucleus accumbens and the paraventricular nucleus of the thalamus^20^. Thus, Hcrt can strongly modulate circuits implicated in OUD.

It has long been noted that human narcoleptics, who have an average 90% loss of Hcrt neurons and very low CSF levels of Hcrt, show little if any evidence of drug abuse, dose escalation or overdose^21^ despite their daily prescribed use of gamma hydroxybutyrate (GHB), methylphenidate and amphetamine. These drugs, which reverse the sleepiness and cataplexy of narcolepsy, are frequently abused in the general population with considerable loss of life^22^-^25^. Human narcoleptics have also been shown to have a greatly reduced reward activation of the VTA, amygdala and accumbens^26^ and altered processing of humor in the hypothalamus and amygdala^27^.

As we discovered, long term self-administration of heroin in humans or giving addictive levels of morphine to mice, produces a substantial increase in the number of detected Hcrt neurons^28^. Cocaine or fentanyl produce similar changes in rats^29^, ^30^. We also reported that heroin self-administration in humans and daily morphine injection in mice for 14 days, produce a marked shrinkage of Hcrt neurons^28^. Because of the role of Hcrt neurons in reward in rodents and pleasure in humans that we^6^, ^12^ and others^17^ have seen, we wanted to rule out the possibility that blocking Hcrt receptors might affect the target regions that mediate rewards and opiate abuse, leading to the findings reported here.

## Results

### Mechanisms underlying the increased number of detected Hcrt neurons and the shrinkage in Hcrt neuron size caused by opiates (Fig 1)

Our prior work in human heroin addicts (Fig 1(a)), and in mice given 50 mg/kg of morphine for 14 or more days, showed that opiates increase the number of Hcrt producing neurons, decrease the size of these neurons, and reduce the intensity of Hcrt neuron immunohistochemical staining^28^. Here we show that these effects are blocked in mice given 50 mg/kg of the opioid antagonist naltrexone 30 min prior to each daily 50 mg/kg morphine dose for 14 days (Fig 1(b),(c)). When naltrexone was given before morphine there were no significant changes in Hcrt cell number (b) (t=0.68, df=6, P=0.53) or size (c) (t=0.21, df=6, P=0.9) (n=4 per condition). **This shows that the morphine effects on Hcrt neurons size and number are mediated by opiate receptors**. But naltrexone not only prevents opiate induced changes in Hcrt neuron number and size and number, it also blocks opiate analgesia^31^.

**Fig 1:**
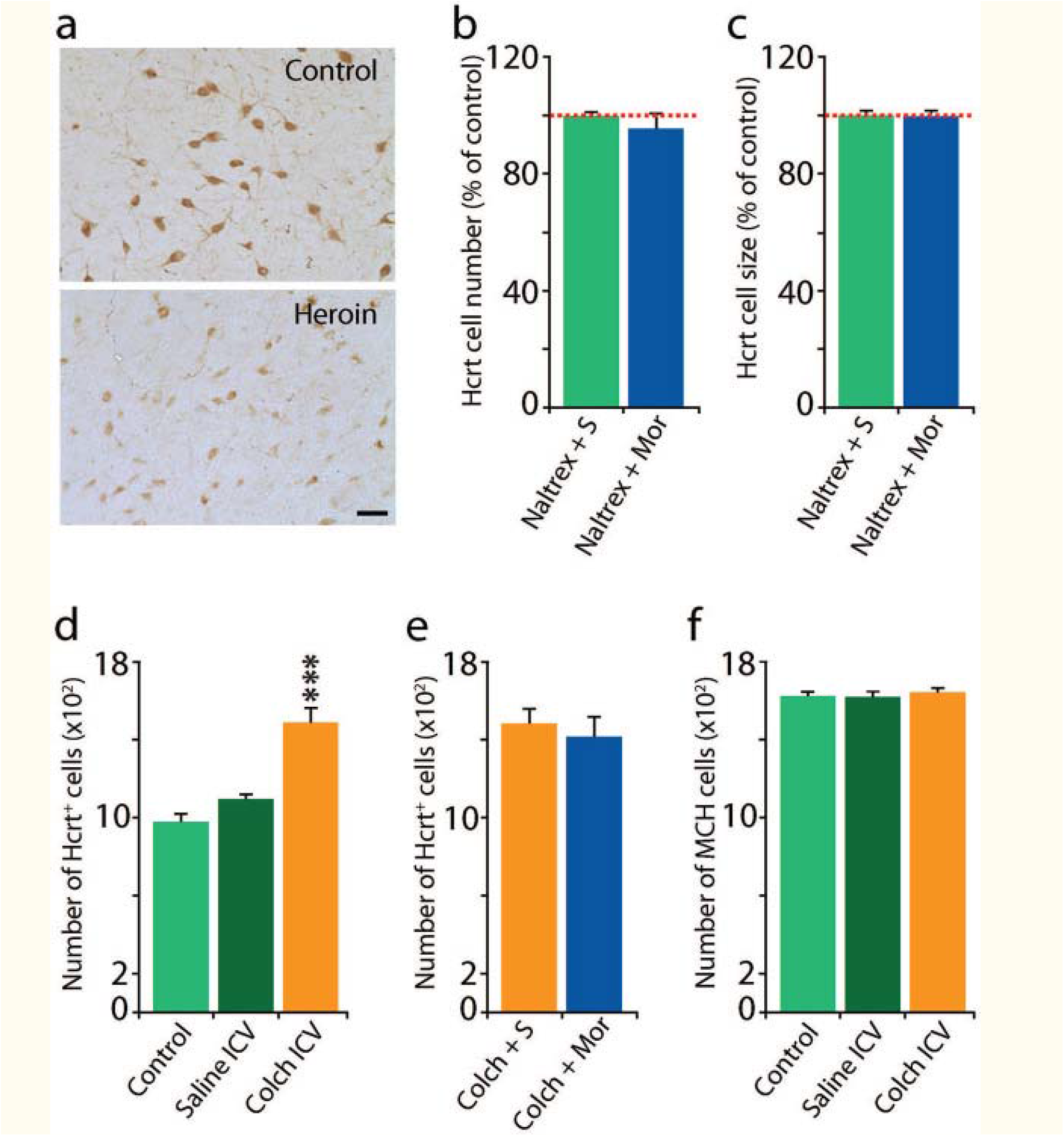
Mechanisms underlying the increased number of detected Hcrt neurons and the shrinkage in Hcrt neuron size caused by opiates^28^. Human heroin addicts (Fig 1(a)), and mice given 50 mg/kg of morphine for a 14 or more days, showed an increase in the number of Hcrt producing neurons and a decreased size of these neurons, with less intense immunohistochemical staining^28^. These effects were blocked in mice by the concurrent administration of the opioid receptor blocker naltrexone Fig 1(b),(c). We observed no difference in Hcrt cell number 1b (n=4/condition, t=0.68, df= 6, P=0.53) or size 1c (t=0.21, df=6, P=0.9) when morphine was preceded by naltrexone. ICV administration of the microtubule transport blocker colchicine (in otherwise drug free mice) increased the number of “detected” Hcrt cells in mice by 44% (Fig 1(d)). The Increase in cell number after colchicine was significant (^***^P<0.001), Tukey post hoc compared to control and saline conditions). (e) Hcrt cell number was not further increased by chronic morphine administration. (f) Colchicine did not significantly alter the number of melanin concentrating hormone (MCH) neurons.

Colchicine, an inhibitor of microtubule polymerization, prevents transport of peptides out of the cell soma. We found that intracerebroventricular injection of colchicine in naïve mice increases the number of Hcrt neurons by 44% Fig 1(d), which is comparable to the percent increase seen in mice after morphine (50 mg/kg for 14 days), i.e. as many as 44% of the neurons capable of producing Hcrt do not produce it at detectable levels under “baseline” conditions in naïve mice (P<0.001, Tukey post hoc, compared to control and saline conditions). Fig 1(e) shows that colchicine together with morphine does not further increase the number of cells labelled, relative to colchicine alone. Together, Fig 1(d) & (e) show that there is a ceiling to morphine effects on Hcrt neuronal number, implying a fixed number of cells capable of producing Hcrt, with 44% beyond the control number of these cells detected in mice and as much as 54% beyond the control number in humans with opiate use disorder^28^. **This is compatible with our conclusion that the morphine induced increase in Hcrt labelled neurons is not due to neurogenesis, but rather due to accumulation of peptide in the cell somas**^28^. Fig 1(f) shows that colchicine does not have any effect on the number of melanin concentrating hormone (MCH) neurons. MCH is a peptide of similar size to Hcrt. MCH neurons are intermingled with Hcrt neurons throughout the hypothalamus.

### Suvorexant also blocks changes in Hcrt cell number and size produced by morphine (Fig 2)

We reported that human heroin addicts^28^ and mice given 50 mg/kg of morphine daily for 14 days have a greatly increased number of detected Hcrt neurons and decreased soma size of Hcrt neurons (Fig 1(a), Fig 2(a),2(b) vehicle (green) vs morphine (blue). Although the dual Hcrt receptor antagonist suvorexant by itself, (yellow) had no significant effect on Hcrt cell number (a) or size (b), blocking Hcrt receptors with suvorexant, 30 mg/kg in 0.5% methylcellulose vehicle, by gavage, 60 min before each daily subcutaneous morphine (50 mg/kg) injection, for 14 days in mice, completely prevented the opioid associated increase in the number (Fig 2a compare blue with red) elicited by morphine. The treatment effect on cell number, vehicle vs morphine, morphine vs suvorexant, and morphine vs suvorexant + morphine was significant, all P=0.0001 Tukey post hoc test. Suvorexant alone did not significantly differ from vehicle alone in effect on cell number (P=0.482).

**Fig 2:**
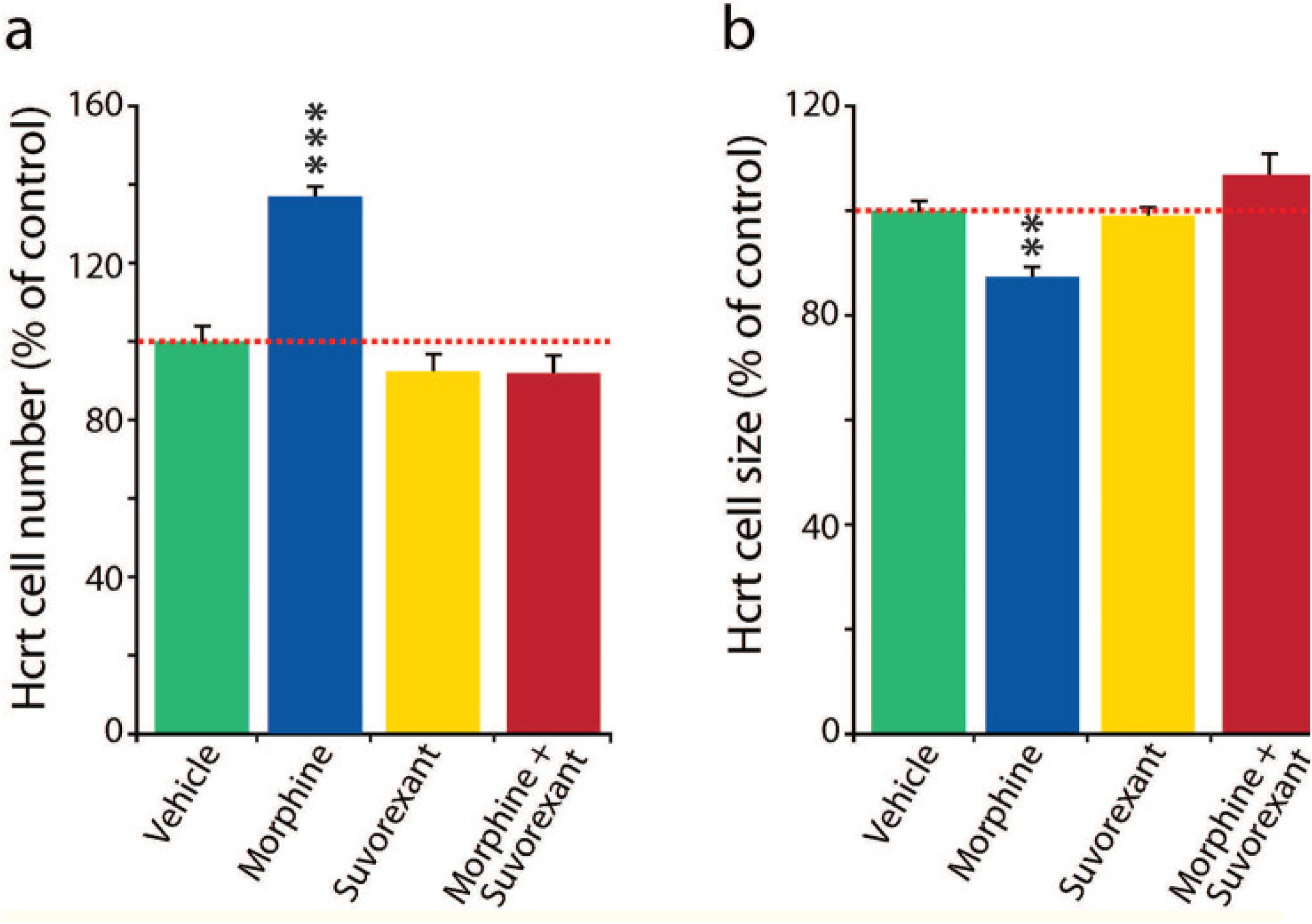
Suvorexant blocked changes in Hcrt cell number and size produced by morphine. Cell number: Fig 2a, Tukey difference test: vehicle-morphine, morphine-suvorexant, and morphine-suvorexant + morphine, all ^***^P=0.0001. Suvorexant alone did not significantly differ from vehicle alone (P=0.482) in effect on cell number. Cell size: Fig 2b, Tukey difference test: vehicle vs morphine P=0.003, morphine vs suvorexant P=0.006, morphine vs suvorexant + morphine ^**^P=0.001. Suvorexant alone did not cause a significant difference from vehicle alone in effect on cell size (P=0.985).

Similarly, suvorexant prevented the reduction in Hcrt soma size produced by morphine (Fig 2b): vehicle vs morphine P =0.003, morphine vs suvorexant P=0.006, morphine vs suvorexant + morphine P=0.001, Tukey post hoc test. Suvorexant alone did not significantly differ from vehicle alone in effect on cell size (P=0.985). **Therefore the Hcrt receptor blocker suvorexant prevents morphine’s effect on Hcrt neuron number and size**.

### Suvorexant blocks morphine induced microglial activation in the hypothalamus and VTA (Fig 3)

Subcutaneous injections of morphine (50 mg/kg) for 14 days increased the number of hypothalamic Iba-1 labeled microglia and increased microglial soma size compared to the saline condition (Fig 3(a),(b)) (P=0.028 and P=0.020,Tukey post hoc, respectively). Suvorexant prevented the morphine induced changes in microglia size and number (Fig 3 (a),(b),(c),(d),(e)). There was no significant difference between microglial size and number in saline, vs suvorexant plus morphine treated mice (P = 0.469 and P=0.958).

**Fig 3:**
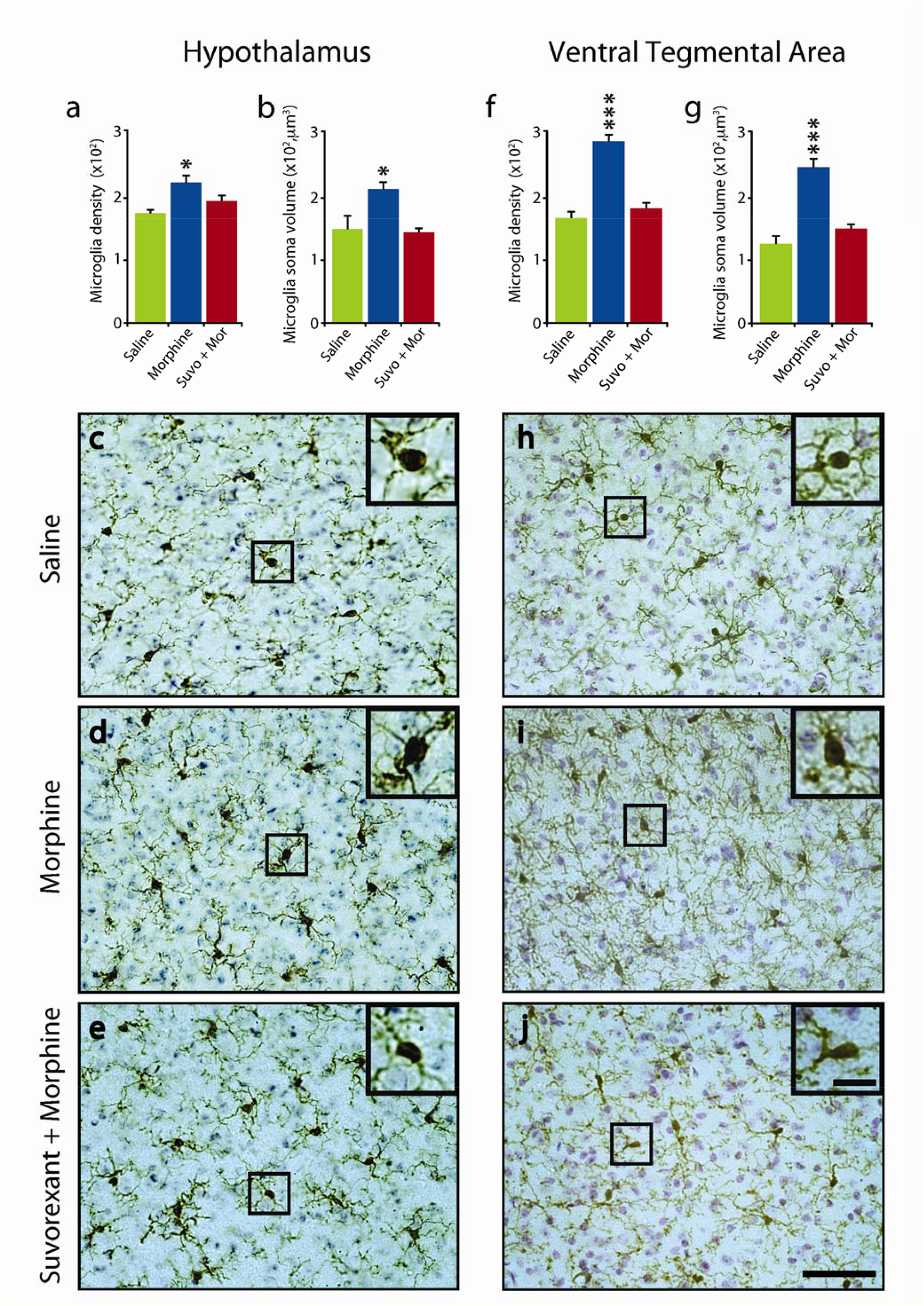
Suvorexant blocked microglial activation in the hypothalamus and VTA. The number (a) and size (b) of microglia in the hypothalamuc was significantly increased by daily morphine (50 mg/kg) treatment (14 d) (n=5 per condition, P=0.028 and P=0.020, Tukey post hoc, respectively) compared to saline treated animals. Administration of suvorexant (30 mg/kg), 60 minutes prior to the morphine treatment completely blocked this effect on microglial number and size (P=0.469 and P=0.958, Tukey post hoc, respectively, relative to saline) (c),(d),(e), show representative images of hypothalamic sections stained for Iba-1 and cresyl violet solution from animals subjected to either saline (c), morphine (d) or suvorexant + morphine (e) illustrating microglial morphology. Similarly, in the VTA region, the number (f) and size (g) of microglia was significantly increased by daily morphine (50 mg/kg) treatment (14d) (n=5 per condition, P=0.001 and P=0.001, Tukey post hoc, respectively) compared to saline treated animals. Administration of suvorexant (30 mg/kg), 60 minutes prior to morphine treatment eliminated this effect in microglial number and size (P=0.137 and P=0.09, Tukey post hoc, respectively, relative to saline). (h),(i),(j), show representative images of section containing the VTA stained for Iba-1 and cresyl violet solution from animals subjected to either saline (h), morphine (i) or suvorexant + morphine (j). Inserts show the area outlined within the black square at higher magnification, illustrating microglial morphology. Scale bar 50μm, insert 10μm ^*^P<0.05, ^***^P<0.001.

Similarly, we observed that morphine treated mice had a significant increase in the number and size of microglial cells in the VTA (Fig 3(f),(g), P=0.001 for both conditions, Tukey post hoc). Suvorexant administration prior to morphine treatment prevented these microglial changes in the VTA. (Fig 3(f),(g),(h),(i),(j)). There was no significant difference between number and size of microglia comparing saline, and morphine + suvorexant in the VTA (P=0.137 and P=0.09 respectively).

### Effect of chronic morphine on Hcrt projections to, and tyrosine hydroxylase (TH) levels in, VTA and substantia nigra (Fig 4)

Daily morphine (50 mg/kg) for 14 days increased Hcrt fluorescence intensity in the VTA Fig 4(a) (P=0.032, n=4/condition, t test). The increase in intensity was accompanied by a significant increase in Hcrt axonal fiber density in the VTA Fig 4(b), P=0.0072, t test), comparable to what we have reported in a prior study in the locus coeruleus (LC)^32^. We found that morphine treatment produces a significant increase in TH immunofluorescence in VTA (Fig 4c,,j),j), P=0.021, t test). We have previously shown that TH immunofluorescence levels have a positive correlation with the amount of TH protein present in the tissue^32^. The increase in TH immunofluorescence in this structure was accompanied by a significant increase in the number of TH+ neurons detected in morphine treated animals compared to controls (Fig 4(d), P=0.0048, t test).

**Fig 4:**
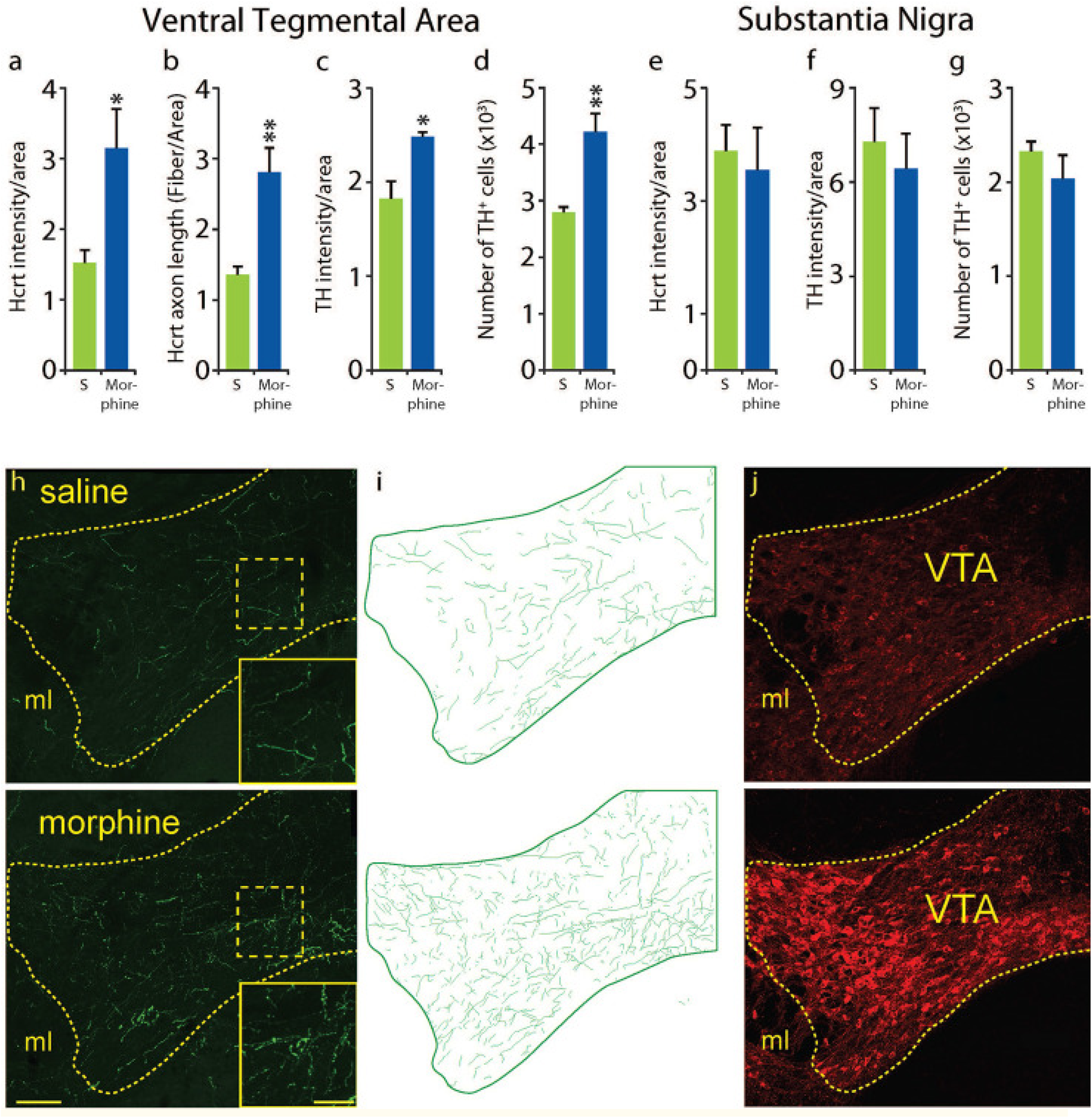
Effect of chronic morphine on Hcrt projections to, and TH levels in, VTA and substantia nigra (SN). Morphine treatment (once a day, 14d, 50 mg/kg, subcutaneous) resulted in a significant increase of Hcrt immunofluorescence intensity per unit of area compared to saline treatment (a) in the VTA (n=4 per condition for all measures, P=0.032, df = 6, t test). This was the result of a significant increase in the total length of Hcrt axons per unit area (b) (P=0.0072, df=6, t test). A significant elevation in TH immunofluorescence intensity per unit of area compared to saline (c) was observed (P=0.021, df=6, t test). This was accompanied by a significant increase in the number of TH+ neurons in VTA (d) (P=0.0048, df=6, t test). Adjacent SN showed no difference in Hcrt (e) P=0.87 or TH (f), P=0.48, immunofluorescence per unit area and no difference in the number of TH + neurons (g), P=0.32) after morphine treatment. The difference in Hcrt innervation is visually apparent (h) between a saline treated (top) and a morphine treated (bottom) animal. Inserts show higher magnification of the square areas in yellow. Hcrt fiber tracings of the VTA (green) (i) in a saline (top) or morphine (bottom) treated animal. Representative sections of the VTA after saline (top) or morphine treatment (bottom) illustrating the difference in TH immunofluorescence (j). Scale bar 100 μm; insert 20 μm. ml = medial lemniscus. ^*^P<0.05,^**^P<0.01.

Although the addiction related structures, LC and VTA, show increased Hcrt innervation and TH immunofluorescence, the motor related substantia nigra (SN), adjacent to VTA had no significant change in Hcrt immunofluorescence intensity after morphine (Fig 4e, P=0.87, t test), TH immunofluorescent intensity (Fig 4f, P=0.48, t test) or TH+ cell number (Fig 4g, P=0.32) after morphine. Fig 4(h),(i),(j) show the difference in Hcrt innervation (green) and TH expression in the VTA (red) of a saline (top) vs a morphine treated animal (bottom).

### Hcrt receptor blockade prevents conditioned morphine anticipation (Fig 5)

Vertical yellow bars indicate light periods in skeleton light-dark cycle. Fig 5a: We studied 3 groups of 6 mice. Drugs were given at ZT(Zeitgeber Time) 4 and ZT5. The first group (blue (V+M) was given vehicle (0.5% methylcellulose) PO at ZT4 and 5 mg/kg of morphine at ZT5. The second group was given suvorexant 30 mg/kg in vehicle (0.5% methylcellulose) at ZT4 and 5 mg/kg of morphine at ZT5 (red (S+M). The third group was given suvorexant 30 mg/kg in vehicle (0.5% methylcellulose) at ZT4 and saline at ZT5 (green (S+S).

**Fig 5:**
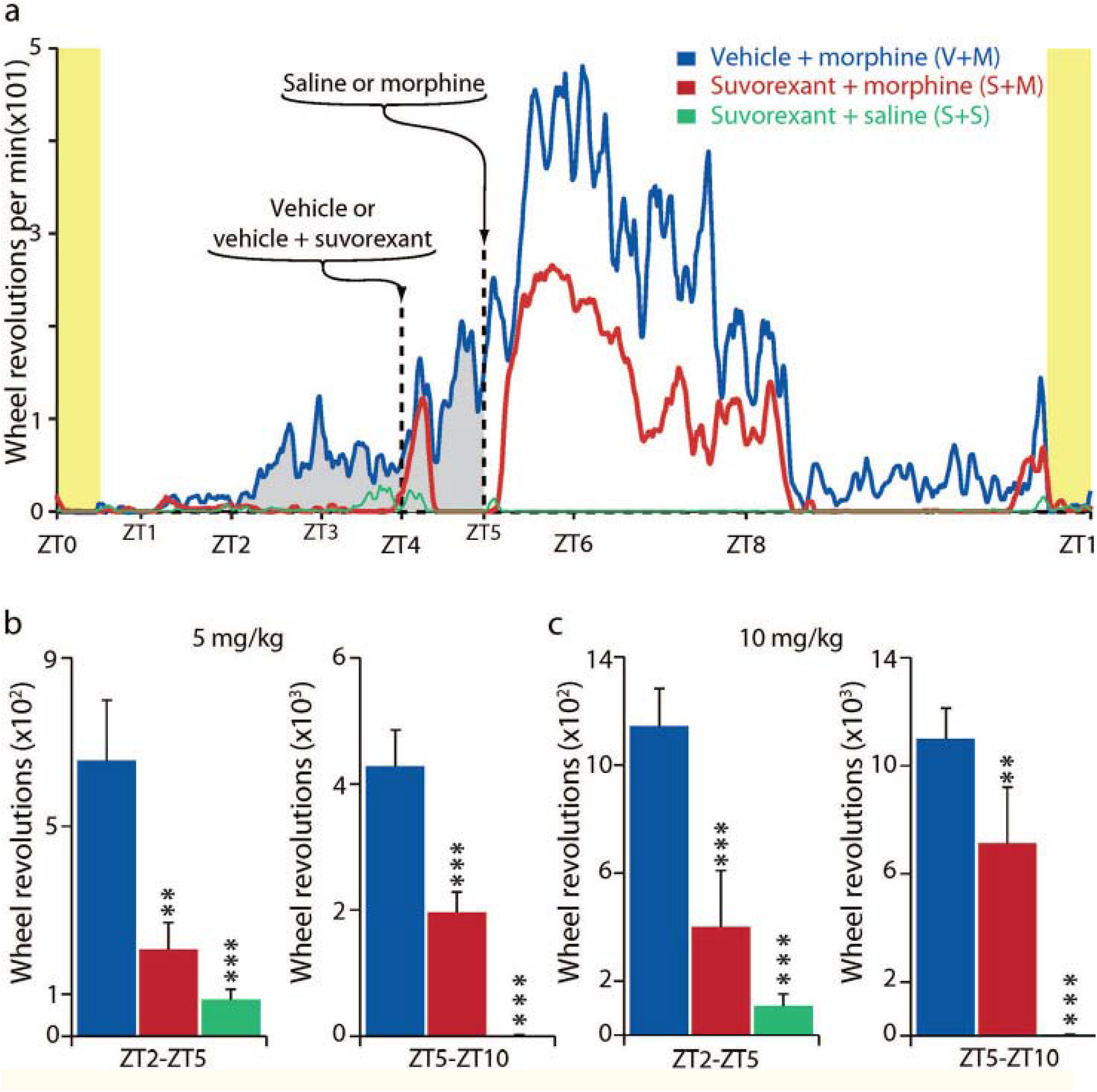
Hcrt receptor blockade prevents conditioned morphine anticipation in morphine treated mice. Fig 5a: Vertical yellow bars on left and right indicate start and end of “skeleton light period.” We studied 3 groups of 6 mice given 5 mg/kg of morphine, and a second cohort of 3 groups of 6 mice given 10 mg/kg of morphine (a total of 36 mice). Groups were given vehicle (0.5% methylcellulose), used for suvorexant, with or without suvorexant (30 mg/kg PO) at ZT4, followed by morphine or saline at ZT5. The 5 mg/kg group is shown in Fig 5a. Anticipatory running (gray fill) is seen in the “vehicle at ZT4 then morphine at ZT5 group” under the (blue line) starting at ZT2 (2 hours after the light on pulse, and 21 hours after the last morphine dose, given on the prior day). The anticipatory running continues to ZT5. Running further increased at ZT5 after morphine injection. The anticipatory running was absent in the group given suvorexant prior to morphine (red line). The group given surorexant followed by saline (green line) also showed no anticipatory running. Fig 5b and andcc indicate total activity during two ZT (ZT2-ZT5 and ZT5-ZT10) interval, for each of the two morphine doses (5 mg/kg (b) and 10 mg/kg (c)). used. ^*^P<0.05, ^**^P<0.01, ^***^ P<0.001, Tukey post hoc test comparing to vehicle + morphine condition.

Fig 5a shows wheel running averaged over the last 12 days of the 14 day study periods for the three morphine 5 mg/kg groups: Anticipatory running (gray fill) is seen in the “vehicle at ZT4 then morphine at ZT5 group” (blue (V+M) line, starting at ZT2 (2 hours after the light on pulse and 3 hours before the daily morphine injection) and continuing to ZT5. Anticipatory wheel running is starting by day 3 presumably as a result of anticipation conditioned by the prior days’ morphine injections at ZT 5, 21 hours earlier^33^-^37^. This anticipation was absent in the suvorexant + morphine group (red (S+M), which received suvorexant 1 hour before the morphine dose 21 hours earlier. Running in both groups further increased at ZT5 after morphine injection. There was also a marked increase in running in both morphine groups’ activity after the mice were handled at ZT4 for vehicle or suvorexant administration. Suvorexant greatly reduced running after morphine injection at ZT5-ZT8 compared to the vehicle-then morphine group, indicating a major dampening by suvorexant on both morphine anticipation and on morphine induced motor excitation. The (green (S+S) line shows the lack of activity in the suvorexant and saline group, which experienced the same handling as the other groups but was not given morphine.

We ran these conditions with both 5 and 10 mg/kg doses of morphine with a virtually identical pattern of activity in both experiments (the 5 mg dose is shown in Fig 5(a)). Bar graphs (Fig 5(b) & Fig 5(c)) indicate total activity during two ZT intervals, for each of the two morphine doses used. Comparisons to vehicle-morphine condition, vehicle vs morphine, morphine vs suvorexant, and morphine vs suvorexant + morphine (P<0.05; P<0.01, P<0.001,Tukey post hoc test, respectively).

### Effect of suvorexant on threshold (Fig 6)

We tested the effect of suvorexant on the pain threshold using an IITC PE34 Incremental Thermal Nociceptive Threshold Analgesia Meter (IITC Life Science Inc). The analgesic effect of morphine on the paw raising response to floor heating can be seen comparing baseline (vehicle alone) to 5 mg/kg or 10 mg/kg, morphine doses. The average analgesic effect (i.e. elevation of nociceptive threshold to heat) (n=6/group, 3 tests) was not significantly diminished by the 30 mg/kg oral dose of suvorexant (actually suvorexant non-significantly increased analgesia) in both 5 mg/kg or 10 mg/kg morphine doses (all P=1.00, Tukey post hoc, comparison between morphine and suvorexant + morphine of either doses). The suvorexant + morphine (5 mg/kg and 10 mg/kg) analgesic effect vs baseline was significant (P<0.01 and P<0.001, respectively, Tukey post-hoc).The suvorexant dosage (30 mg/kg) that preserved opiate analgesia is the same as that which completely prevented the chronic morphine associated increase in Hcrt cell number and decrease in size seen in Fig 2, the microglial activation seen in Fig 3 and the morphine anticipation seen in Fig 5.

**Fig 6:**
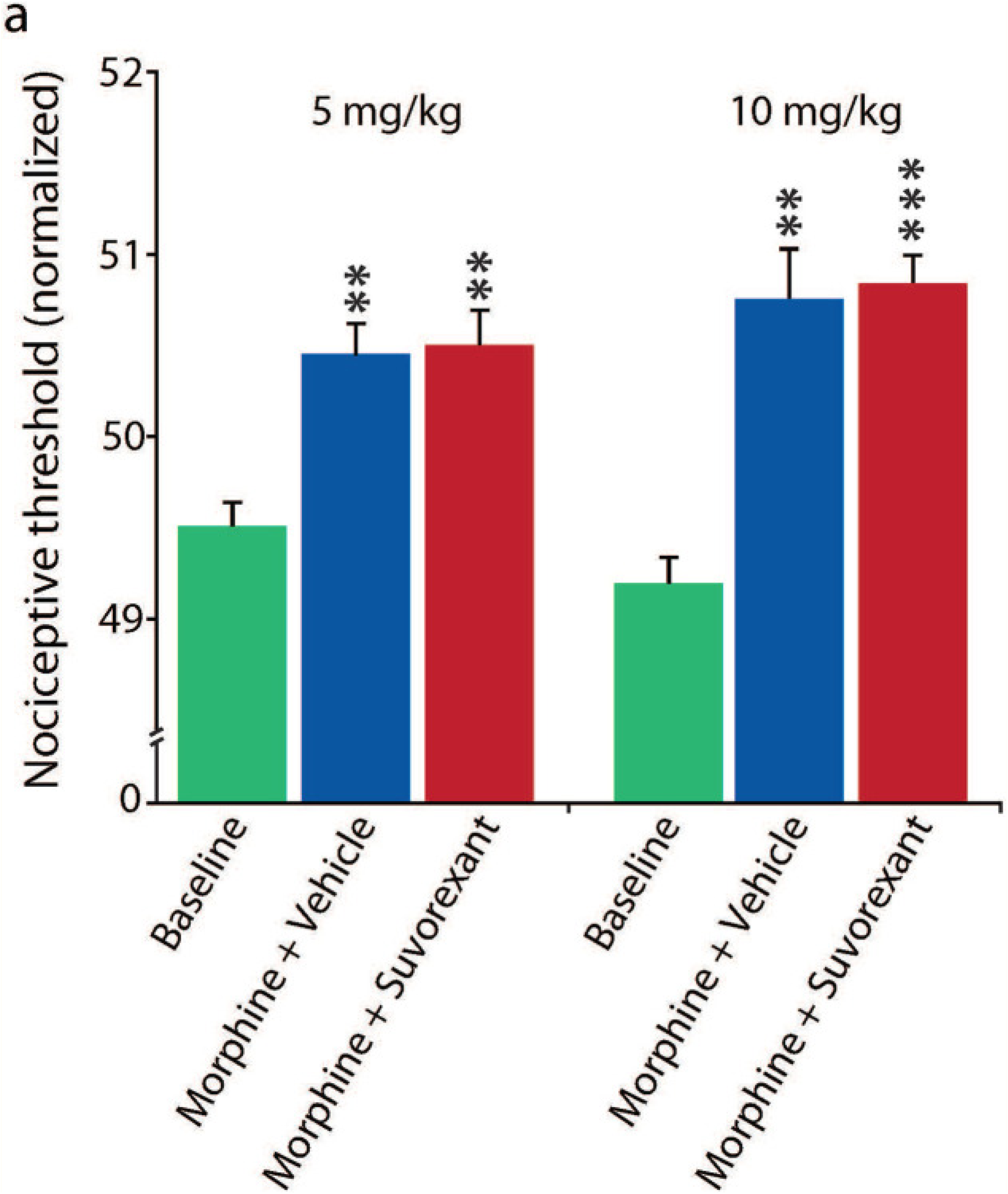
Effect of suvorexant on pain. The analgesic effect of morphine on the paw raising response to floor heating can be seen comparing baseline (vehicle alone) to 5 mg/kg or 10 mg/kg, morphine doses. The average analgesic effect (i.e. elevation of nociceptive threshold to heat) (n=6/group, 3 tests) was not significantly diminished by 30 mg/kg oral (by gavage) dose of suvorexant (actually suvorexant non-significantly increased analgesia) in both 5 mg/kg or 10 mg/kg morphine doses (all P=1.00, Tukey post hoc comparisons between morphine + vehicle and suvorexant + morphine of either doses). The suvorexant + morphine (5 mg/kg and 10 mg/kg) analgesic effect vs baseline was significant (P<0.01 and P<0.001, respectively, Tukey post hoc). The suvorexant dosage (30 mg/kg) that completely preserved opiate analgesia (Fig 6) is the same as that which prevented the chronic morphine associated increase in Hcrt cell number and decrease in size seen in Fig 2, the microglial activation seen in Fig 3 and the morphine anticipation seen in Fig 5. ^**^P<0.01 and ^**^P<0.001.

## Discussion

As shown in^28^ and in Fig 1(a), long-term self-administration of heroin in humans or administration of addictive levels of morphine to mice, as well as administration of cocaine or fentanyl to rats^29 30^ produce an increase in the number and decrease in size of detected Hcrt neurons. We now show that administration of the opioid antagonist naltrexone prevents these morphine induced changes in size and number of Hcrt neurons, confirming that opiate receptor activation is required for this effect.

Colchicine, an inhibitor of microtubule polymerization that prevents transport of peptides out of the cell soma^38^, produces an increase in the number of detectable Hcrt neurons, likely due to the accumulation of Hcrt in the cell soma (Fig 1). The percentage increase in the number of detected Hcrt neurons after colchicine is comparable in magnitude to the increase in the number of Hcrt neurons produced by chronic administration of heroin in humans or morphine in naïve mice or rats^28^, ^29^, suggesting that the opiate induced increase in the number of neurons staining for Hcrt results from increased accumulation of Hcrt in neurons that have sub-detection levels of Hcrt under baseline conditions, rather than being caused by neurogenesis of “new” Hcrt neurons^28^. Administration of both morphine and colchicine does not increase the number of Hcrt neurons beyond that produced by colchicine alone, consistent with this conclusion^28,39^.

We have reported that morphine increases Hcrt neuronal projections to the locus coeruleus^6^. We now show in Fig 4, that 50 mg/kg of morphine for 14 days also increases projections to the VTA, a region that has been strongly implicated in opiate addiction^40^-^42^. We find that Hcrt projections to the substantia nigra, a TH cell containing region adjacent to VTA but not generally implicated in opiate addiction, are not significantly increased by opiates, indicating a local regulation of Hcrt projections.

It has been observed that opiates produce activation of microglial cells throughout the brain. Microglia have both opioid^43^-^45^ and Hcrt^46^ receptors. We now report that suvorexant, a dual Hcrt receptor antagonist, prevents morphine induced activation of microglia in the hypothalamus and in the ventral tegmental area (Fig 3), showing a key role of Hcrt in opiate induced microglial proliferation.

An *in vitro slice* study found that opioids decrease the activity of Hcrt neurons^47^. However our in vivo data^28^ shows that systemic administration of morphine greatly increases the activity of Hcrt neurons and the brain level of Hcrt^28^. The effects of opioid agonists can be exerted not only in plasma membrane receptors and endosomes, but also in the Golgi apparatus^48^, suggesting possible intracellular pathways mediating the opioid induced decrease of Hcrt neuronal size that we have reported^28^.

Narcoleptic humans, who have an average 90% loss of Hcrt producing neurons^49^-^51^, are resistant to addiction^21^. Removal of Hcrt neurons in mice significantly reduces aversion elicited by naloxone precipitated morphine withdrawal^32^. Because of the role of Hcrt neurons in reward in rodents and pleasure in humans that we^6^, ^12^ and others^17^, ^52^, ^53^ have seen, we wanted to rule out the possibility that blocking Hcrt receptors might also affect the opiate elicited shrinkage and increase in the number of detectable Hcrt neurons that we discovered^28^. We thought that this blockade might have no effect, since we assumed that the morphine induced changes in Hcrt neurons were solely the result of direct opioid action on mu opioid receptors on Hcrt neurons (Fig 1). However, to our surprise, we found that administration of the dual Hcrt receptor antagonist completely prevented the increase in number and the decrease in size of Hcrt neurons produced by opiate administration (Fig 2). This finding led us to further test the effect of suvorexant on morphine anticipation and on analgesia.

We utilized a wheel running model of addictive anticipation of drug administration^36^. After a few days of daily injection of morphine at ZT5, mice anticipated these injections by vigorous running starting at ZT2, at morphine doses as low as 5 mg/kg (Fig 5). However when morphine injections were preceded by suvorexant administration no anticipatory wheel running occurred. Note that the anticipation was occurring 22 hours after the prior suvorexant injection 21 hours after the prior morphine injection and 3 hours before the next morphine injection, at a time when minimal amounts of morphine or suvorexant would still be present in the mice. It seems likely that it was the presence of suvorexant when the morphine was injected that prevented the condtioned anticipatory running 22 hours later. The suvorexant effect on morphine anticipation may be a result of the blockade of self-excitation of the Hcrt neuronal population^54,55^ This blockade of Hcrt receptors appears to provide the same protection against substance use disorder that is present in the absence of Hcrt neurons in human and animal narcoleptics and after naloxone administration (Fig 1).

At the suvorexant dose used, even though given in the light period (the normal sleep period) no sleep occurred in either the mice given morphine or the mice given suvorexant + morphine, as evidenced by wheel running and video observation. This is due to the well-known arousing quality of opioids in mice and rats^56^. Therefore, even though suvorexant is a “sleeping pill” at certain doses in humans (the dual Hcrt receptor antagonist suvorexant is marketed as Belsomra, with newer dual Hcrt receptor antagonists daridorexant in Quviviq and lemborexant in Dayvigo), sleep did not mediate the observed effect of this drug on morphine induced changes in Hcrt cell number and size shown in Figs 1&2. It is useful to recall that morphine itself is soporific in humans, which is why morphine was named after Morpheus. Sleepiness does not prevent the opiate addictive process^57^.

Suvorexant prevents the changes in Hcrt neuron number and size, produced by chronic administration of opiates (Fig 2), the microglial activation seen in Fig 3, and the anticipatory motor activation in mice expecting opioid administration (Fig 5). However, we find that opiate analgesia is not at all diminished by suvorexant (Fig 6). These findings suggest that administration of Hcrt receptor antagonists combined with opioids for pain relief in humans may greatly reduce addiction risk and consequent morbidity while providing maximal analgesia.

## Methods

### Animal Usage

All procedures were approved by the Institutional Animal Care and Use Committees of the University of California at Los Angeles and of the Veterans Administration Greater Los Angeles Health Care System. Experiments were performed in C57BL/6 mice. A total of 151 male mice were used in the current study. All experimental procedures were started when animals reached 3 months of age. Animals were kept in a room maintained at 22±1°C on a 12 h light (135 lux) dark (0.03 lux) cycle (lights on at 7 AM and off at 7 PM) or on a “skeleton” light schedule as described below.

### Drugs

**Morphine sulfate** (Hospira Inc., Lake Forest, IL, USA) and **Naltrexone hydrochloride**, (Sigma-Aldrich, N3136, Lot # SLBF858548, Saint Louis, MO, USA) 50 mg/kg were used. Morphine and naltrexone were dissolved in sterile saline immediately before subcutaneous administration. **Suvorexant** (Belsomra, Merck NJ, USA) was suspended in 0.5% methylcellulose in water and orally administered by gavage.

### Tissue processing

Animals were anesthetized intraperitoneally (IP) with Fatal-Plus pentobarbital solution (150 mg/kg) and then perfused transcardially with PBS (0.1M, pH 7.4), followed by 4% formaldehyde in PBS. Brains were removed and post-fixed for 72 hours in 4% formaldehyde in PBS, followed by 20% sucrose in PBS for 24 hours and 30% for 48 hours. Brains were frozen and cut into 40 μm coronal sections using a sliding microtome (American Optical, USA). The sections were sorted into one in three-section compartments. Immunohistochemical procedures were performed immediately. The remaining tissue was transferred to a cryoprotectant solution and stored at −20°C. Mice that we compared were always sacrificed and processed together. Perfusion, histology and microscopic analysis were performed by investigators blind to the procedures performed prior to sacrifice.

### Immunostaining for brightfield microscopy

All immunohistochemical procedures were performed by sequential incubation of free-floating sections. For detection of Hcrt and MCH, sections were first incubated for 30 min in 0.5% H_2_O_2_ in PBS to block endogenous peroxidase activity. After thorough washing with PBS, the sections were placed for 2 hours in 1.5% normal goat serum (NGS) in PBS containing 0.25% Triton X (PBST) and incubated for 72 hours at 4°C in a PBST solution containing rabbit anti-Hcrt-1 primary antibody (1:10000, H-003-30, Lot # 01108, Phoenix Pharmaceuticals Inc.) or rabbit anti-MCH (1:20000, H-070-47, Lot # 01629-5, Phoenix Pharmaceuticals Inc.), followed by the corresponding biotinylated secondary antibody (1:400, Vector Laboratories)in PBST for 2 h, and avidin-biotin-peroxidase complex (1:300, ABC Elite Kit, Vector Laboratories) in PBS for 2 h. The tissue-bound peroxidase was then developed using the diaminobenzidine tetrahydrochloride (DAB) method, which consisted of tissue immersion in 0.02% DAB and 0.03% hydrogen peroxide in 10 ml PBS. Microglia were identified using the canonical marker, ionized calcium binding adaptor molecule-1 (Iba-1)^58^. Prior to Iba-1 immunostaining, an antigen retrieval procedure was performed by incubating the sections in 10mM sodium citrate (pH 8.5) at 80°C for 30 min. The sections were then cooled to room temperature in sodium citrate, washed with PBS and following the same staining procedures used to detect Hcrt-1 using primary antibody goat anti-Iba-1 (ab5076, Abcam, Lot#GR3403958, 1:10000) and normal rabbit serum.

We previously standardized our DAB method and established an 8-minute optimal developing time and used this precise duration in all of our studies. All developing solutions were prepared in one container, homogenized and aliquoted to the respective developing wells. Developing procedures were performed with room lights off and the wells containing tissue were wrapped with aluminum foil to protect them from light exposure. Wells were agitated at 55 rpm. Developing solutions were used only once.

### Brightfield microscopy

The number, distribution and size of Hcrt+ and Ibal-1+ cells and the number and distribution of MCH were assessed using a Nikon Eclipse 80i microscope with three axis motorized stage, video camera, Neurolucida interface and Stereoinvestigator software (MicroBrightField Corp.). Cell counting was performed bilaterally using either 40x or 60x objective and cell size was determined using the Neurolucida Nucleator probe. Iba-1+ quantification was performed bilaterally in the middle of the Hcrt neuronal field by placing a square (250 μm x 250 μm) dorsal to the fornix, with the lower corners of the square equidistant from the center of the fornix. Quantification of Iba-1+ cells was performed bilaterally in the same manner in the VTA, by placing a square (250 μm x 250 μm) medial to the medial lemniscus. All counting and cell measurements were performed by a trained histologist, always blind to the experimental condition. In every case, the same individual counted both the experimental and control tissue. Only neurons with an identifiable nucleus were counted. In addition to quantitative assessments, we look for nuclear fragmentation,chromatolysis, inclusions, varicosities and other abnormalities^7^,^12^, ^32^, ^39^, but we did not see these phenomena in the current study.

### Immunostaining for confocal microscopy

For identification of dopamine (DA) containing neurons in the VTA and substantia nigra (SN) regions we used the immunohistochemical detection of TH enzyme. The sections were first incubated in PBST containing 1.5% of both NGS and normal donkey serum (NDS), followed by co-incubation with primary antibodies rabbit anti-Hcrt-1 (H-003-36, Phoenix Pharmaceuticals, USA, 1:2000, Lot # 01108) and sheep anti-TH (ab113, Abcam, USA, 1:1000, Lot # GR 3277795-15) overnight at room temperature in PBST. Next, we washed the sections and incubated them in PBST containing the corresponding secondary antibody tagged with fluorophores that match our microscope filters (1:300, Alexafluor 488 goat anti-rabbit, A11008, Lot # 2557379, Alexafluor 555 donkey anti-sheep, A 21436, Lot # 2420712 ThermoFisher Scientific, US), 1% NGS and 1% NDS, with lights off and samples wrapped in aluminum foil. Tissue was mounted and cover slipped using Vector Shield anti fade mounting media (H 1000, Vector Laboratories, Burlingame, CA, USA, Lot # 2E00806). All tissue sections from experimental and control animals were stained at the same time and with the same antibody lot. To quantify the number of TH+ neurons, the same mounting media containing 42, 6-diamidino-2-phenylindole (DAPI) was used (H 1200, Vector Laboratories, Burlingame, CA, USA, Lot # 2E0815). All tissue sections from experimental and control animals were stained at the same time and with the same antibody lot.

### Confocal microscopy

The number and distribution of Hcrt fibers and TH cell bodies was assessed using a Zeiss LSM 900 (Imager Z2 AX10, Jena, Germany) confocal microscope equipped with the appropriate lasers. Every section that contained the VTA and adjacent SN was imaged at 1 μm optical planes; 28 ± 1.4 optical planes were obtained per section respectively. Quantification was performed bilaterally on every third section throughout the region of interest. Immunofluorescence intensities and area measurements were obtained using the Zeiss proprietary software ZEN^®^. The area was defined by the size of the region containing TH neuronal bodies in each structure. Total immunofluorescence for TH and Hcrt was divided by the corresponding area, and bilateral areas in the same section were averaged. Cell number and fiber distribution were determined using the Adobe Illustrator program. Every stack of images was loaded in this program such that each optical plane was placed in a distinct layer and individual TH containing neurons were identified and marked. Using this method of quantification eliminates double counting of cells, which is critical in the analysis of structures with a high density of neuronal bodies like the VTA and SN.

For the analysis of Hcrt fiber length and distribution, the middle optical plane was chosen for matching sections that contained the VTA. Individual fibers were drawn and measured and the total fiber length was then calculated based on the area used to calculate the immunofluorescence intensities.

### Colchicine procedure

Two groups of animals (n=4 per group) were subjected to intracerebroventricular (ICV) injection of either saline solution or saline solution containing 20 μg/μl of colchicine, a microtubule disruptor that prevents peptide transport and increases levels of neuropeptides in cell somas^38^. Another two groups of animals (n=4 per group) received 14 d of either saline or morphine (50mg/kg) before ICV injections of colchicine. An additional 4 naïve animals served as the control. Anesthesia was induced with a mixture of ketamine/xylazine (100 mg/kg/15 mg/kg, i.p.) and then maintained with a gas mixture of isoflurane in oxygen (1-3%) after the animals were placed in the stereotaxic device. Body temperature was maintained with a water-circulating heating pad (Gaymar Industries, Orchard Park, NY, USA). The head was positioned in a stereotaxic frame and the skull was exposed. A hole was drilled at coordinates corresponding to the lateral ventricle (AP: −0.5 mm, L: −1 mm, relative to bregma). A Hamilton microsyringe was lowered until the ventricle was reached (H-2.8 mm, relative to the skull surface). Infusion was made in increments of 0.2 μl every 10 minutes for 40 minutes to obtain a final volume of 1 μl. The needle was held in place for another 10 minutes before being slowly withdrawn. Ventricular localization was confirmed by observing free flowing cerebrospinal fluid after the withdrawal of the needle. A small piece of sterile bone wax was placed over the hole and the skin sutured. All subjects recovered from the anesthesia within 30 minutes after the end of the procedure. Animals were carefully monitored and sacrificed 52 h later between ZT 13 and ZT 15 for immunohistochemical procedures.

### Morphine anticipation

To measure conditioned addictive anticipation, Juarez-Portilla et al.^36^ developed a wheel running test in mice. This technique has been used to quantify drug anticipation and appetitive changes^37^. We adapted this method for measuring morphine anticipation. The running wheels were low-profile running wheels for mice, placed inside the testing cages, L 48.3 cm, W 26.7 cm and H 40.6 cm and wirelessly monitored (Med Associates, Model ENV-047). Mice were first acclimated to a “skeleton light cycle” (lights on at 6:00 AM and off at 6:00 PM, 60 min total light per 24 h period, light was on for 30 min at the beginning of the “light” period and 30 min at the end of the 12 h “light” cycle) for 4 weeks. This lighting schedule disinhibits running wheel behavior during a period in which light would otherwise inhibit wheel running, then running wheels were introduced for an additional 14 days before the start of drug administration. Running wheel activity was collected and analyzed using Med Associates software (SOF-861). Continuous video monitoring and the wheel running record showed that no sleep occurred for more than 4 h after morphine administration in any mouse.

A total of 36 male mice received 14 days of orally administered suvorexant (30 mg/kg) or vehicle (0.5% methylcellulose) at 10:00 AM (ZT4) followed by morphine (5 or 10 mg/kg, subcutaneous) or saline (0.05 ml, subcutaneous) at 11:00 AM (ZT5).

### Analgesia measurement

We measured the effect of morphine with and without suvorexant on the pain threshold using an IITC PE34 Incremental Thermal Nociceptive Threshold Analgesia Meter (IITC Life Science Inc), which raises the temperature of its aluminum surface at 6°C/min in each trial. When the mouse licked or shook a hindlimb or jumped, the experimenter immediately removed it from the apparatus and pressed the stop switch on the apparatus to record the surface temperature. Baseline threshold was established on 3 consecutive days, with 3 tests/day. Then, two tests, a non-drug pre-test and a 60 min post-drug, were done daily. Thirty six male mice (n=6 per group) were used. Suvorexant (30 mg/kg in 0.5% methylcellulose) or vehicle (0.5% methylcellulose) was given orally 60 min before morphine (5 mg/kg or 10 mg/kg, subcutaneous). The threshold test started 60 min after morphine. The animal was checked for any skin inflammation or lesion from the thermal test and would have been removed immediately from the experiment for treatment if either occurred, but this did not happen.

## Statistical analysis

Data were subjected to ANOVA followed by Tukey post hoc test comparisons or t test. All such tests were two-tailed. Results were considered statistically significant if P<0.05. The number of subjects in each experimental procedure is indicated by degrees of freedom (df).

## Acknowledgements

Support: DA034748, HL148574, Medical Research Service of the Department of Veterans Affairs.

## Reference List

1. Cicero T.J. No end in sight: The abuse of prescription narcotics. Cerebrum. cer–11-15 (2015). [PMC free article] [PubMed] [Google Scholar]

2. Mack K.A., Jones C.M., & McClure R.J. Physician dispensing of oxycodone and other commonly used opioids, 2000-2015, United States. Pain Med. 19, 990–996 (2018). [PMC free article] [PubMed] [Google Scholar]

3. Bass C. & Yates G. Complex regional pain syndrome type 1 in the medico-legal setting: High rates of somatoform disorders, opiate use and diagnostic uncertainty. Med Sci Law. 58, 147–155 (2018). [PubMed] [Google Scholar]

4. Kelly M.M., Reilly E., Quinones T., Desai N., & Rosenheck R. Long-acting intramuscular naltrexone for opioid use disorder: Utilization and association with multi-morbidity nationally in the Veterans Health Administration. Drug Alcohol Depend. 111–117 (2018). [PubMed] [Google Scholar]

5. Parthvi R., Agrawal A., Khanijo S., Tsegaye A., & Talwar A. Acute opiate overdose: an update on management strategies in emergency department and critical care unit. Am J Ther. 26, e380–e387 (2019). [PubMed] [Google Scholar]

6. McGregor R., Wu M.-F., Barber G., Ramanathan L., & Siegel J.M. Highly specific role of hypocretin (orexin) neurons: differential activation as a function of diurnal phase, operant reinforcement vs. operant avoidance and light level. Journal of Neuroscience 31, 15455–15467 (2011). [PMC free article] [PubMed] [Google Scholar]

7. Wu M.F., Nienhuis R., Maidment N., Lam H.A., & Siegel J.M. Role of the hypocretin (orexin) receptor 2 (Hcrt-r2) in the regulation of hypocretin level and cataplexy. J Neurosci. 31, 6305–6310 (2011). [PMC free article] [PubMed] [Google Scholar]

8. Wu M.F., Nienhuis R., Maidment N., Lam H.A., & Siegel J.M. Cerebrospinal fluid hypocretin (orexin) levels are elevated by play but are not raised by exercise and its associated heart rate, blood pressure, respiration or body temperature changes. Arch. ital. Biol. 149, 492–498 (2011). [PMC free article] [PubMed] [Google Scholar]

9. Mileykovskiy B.Y., Kiyashchenko L.I., & Siegel J.M. Behavioral correlates of activity in identified hypocretin/orexin neurons. Neuron 46, 787–798 (2005). [PMC free article] [PubMed] [Google Scholar]

10. Kiyashchenko L.I. et al. Release of hypocretin (orexin) during waking and sleep states. J Neurosci 22, 5282–5286 (2002). [PMC free article] [PubMed] [Google Scholar]

11. James M.H., Fragale J.E., O’Connor S.L., Zimmer B.A., & Aston-Jones G. The orexin (hypocretin) neuropeptide system is a target for novel therapeutics to treat cocaine use disorder with alcohol co-abuse. Neuropharmacology 108359 (2020). [PMC free article] [PubMed] [Google Scholar]

12. Blouin A.M. et al. Human hypocretin and melanin-concentrating hormone levels are linked to emotion and social interaction. Nature Communications 4:1547, 1547 (2013). [PMC free article] [PubMed] [Google Scholar]

13. Farahimanesh S., Zarrabian S., & Haghparast A. Role of orexin receptors in the ventral tegmental area on acquisition and expression of morphine-induced conditioned place preference in the rats. Neuropeptides. 66, 45–51 (2017). [PubMed] [Google Scholar]

14. Meye F.J., van Zessen R., Smidt M.P., Adan R.A.H., & Ramakers G.M.J. Morphine withdrawal enhances constitutive opioid receptor activity in the ventral tegmental area. Journal of Neuroscience 32, 16120–16128 (2012). [PMC free article] [PubMed] [Google Scholar]

15. Sarti F., Borgland S.L., Kharazia V.N., & Bonci A. Acute cocaine exposure alters spine density and long-term potentiation in the ventral tegmental area. Eur J Neurosci. 26, 749–756 (2002). [PubMed] [Google Scholar]

16. Stefano G.B. & Kream R.M. Endogenous morphine synthetic pathway preceded and gave rise to catecholamine synthesis in evolution (Review). Int J Mol. Med. 20, 837–841 (2007). [PubMed] [Google Scholar]

17. Baimel C. et al. Orexin/hypocretin role in reward: implications for opioid and other addictions. Br J Pharmacol 172, 334–348 (2015). [PMC free article] [PubMed] [Google Scholar]

18. Narita M. et al. Direct involvement of orexinergic systems in the activation of the mesolimbic dopamine pathway and related behaviors induced by morphine. J Neurosci. 26, 398–405 (2006). [PMC free article] [PubMed] [Google Scholar]

19. Vittoz N.M., Schmeichel B., & Berridge C.W. Hypocretin /orexin preferentially activates caudomedial ventral tegmental area dopamine neurons. The European journal of neuroscience 28, 1629–1640 (2008). [PMC free article] [PubMed] [Google Scholar]

20. Peyron C. et al. Neurons containing hypocretin (orexin) project to multiple neuronal systems. J Neurosci 18, 9996–10015 (1998). [PMC free article] [PubMed] [Google Scholar]

21. Guilleminault C. & Cao M.T. Narcolepsy: Diagnosis and management in Principles and Practice of Sleep Medicine (eds. Kryger M.H., Roth T. & Dement W.C.) 957–968 (Elsevier Saunders, Missouri, 2011). [Google Scholar]

22. Galloway G.P. et al. Gamma-hydroxybutyrate: an emerging drug of abuse that causes physical dependence. Addiction. 92, 89–96 (1997). [PubMed] [Google Scholar]

23. Perrotti L.I. et al. Distinct patterns of DeltaFosB induction in brain by drugs of abuse. Synapse. 62, 358–369 (2008). [PMC free article] [PubMed] [Google Scholar]

24. Zhu J., Spencer T.J., Liu-Chen L.Y., Biederman J., & Bhide P.G. Methylphenidate and opioid receptor interactions: A pharmacological target for prevention of stimulant abuse. Neuropharmacology 61, 283–292 (2011). [PMC free article] [PubMed] [Google Scholar]

25. Darke S., Peacock A., Duflou J., Farrell M., & Lappin J. Characteristics and circumstances of death related to gamma hydroxybutyrate (GHB). Clinical Toxicology 58, 1028–1033 (2020). [PubMed] [Google Scholar]

26. Ponz A. et al. Abnormal activity in reward brain circuits in human narcolepsy with cataplexy. Ann Neurol. 67, 190–200 (2010). [PubMed] [Google Scholar]

27. Schwartz S. et al. Abnormal activity in hypothalamus and amygdala during humour processing in human narcolepsy with cataplexy. Brain 131, 514–522 (2007). [PubMed] [Google Scholar]

28. Thannickal T.C. et al. Opiates increase the number of hypocretin-producing cells in mouse and human brain, and reverse cataplexy in a mouse model of narcolepsy. Sci Transl Med 10, pii: eaao4953-doi: 10.1126/scitranslmed.aao4953. (2018). [PMC free article] [PubMed] [CrossRef] [Google Scholar]

29. Fragale J.E., James M.H., & Aston-Jones G. Intermittent self-administration of fentanyl induces a multifaceted addiction state associated with persistent changes in the orexin system. Addict Biol 2020/August/14, e12946 (2021). [PMC free article] [PubMed] [Google Scholar]

30. James M.H. et al. Increased number and activity of a lateral subpopulation of hypothalamic orexin/hypocretin neurons underlies the expression of an addicted state in rats. Biol Psychiatry doi: 10.1016/j.biopsych. (2019). [PMC free article] [PubMed] [CrossRef] [Google Scholar]

31. Vickers A.P. Naltrexone and problems in pain management. BMJ 332, 132–133 (2006). [PMC free article] [PubMed] [Google Scholar]

32. McGregor R. et al. Hypocretin/orexin interactions with norepinephrine contribute to the opiate withdrawal syndrome. Journal of Neuroscience 42, 255–263 (2022). [PMC free article] [PubMed] [Google Scholar]

33. Esmaili-Shahzade-Ali-Akbari P., Hosseinzadeh H., & Mehri S. Effect of suvorexant on morphine tolerance and dependence in mice: Role of NMDA, AMPA, ERK and CREB proteins. NeuroToxicology 84, 64–72 (2021). [PubMed] [Google Scholar]

34. Gillman A.G., Leffel J.K., Kosobud A.E.K., & Timberlake W. Fentanyl, but not haloperidol, entrains persisting circadian activity episodes when administered at 24- and 31-h intervals. Am J Psychiatry 2004. Nov. ;161. (11):2126.–8. 2009/07/10, 102–114 (2009). [PMC free article] [PubMed] [Google Scholar]

35. Gillman A.G., Rebec G.V., Pecoraro N.C., & Kosobud A.E.K. Circadian entrainment by food and drugs of abuse. Behav Processes 2019/May/24, 23–28 (2019). [PMC free article] [PubMed] [Google Scholar]

36. Juarez-Portilla C. et al. Brain activity during methamphetamine anticipation in a non-invasive self-administration paradigm in mice. eNeuro 5, ENEURO (2018). [PMC free article] [PubMed] [Google Scholar]

37. LeSauter J., Balsam P.D., Simpson E.H., & Silver R. Overexpression of striatal D2 receptors reduces motivation thereby decreasing food anticipatory activity. The European journal of neuroscience 51, 71–81 (2020). [PMC free article] [PubMed] [Google Scholar]

38. Correa L.M., Nakai M., Strandgaard C.S., Hess R.A., & Miller M.G. Microtubules of the mouse testis exhibit differential sensitivity to the microtubule disruptors carbendazim and colchicine. Toxicological Sciences 69, 175–182 (2002). [PubMed] [Google Scholar]

39. McGregor R., Shan L., Wu M.F., & Siegel J.M. Diurnal fluctuation in the number of hypocretin/orexin and histamine producing: Implication for understanding and treating neuronal loss. PLoS ONE 12, (2017). [PMC free article] [PubMed] [Google Scholar]

40. Thomas T.S., Baimel C., & Borgland S.L. Opioid and hypocretin neuromodulation of ventral tegmental area neuronal subpopulations. Br J Pharmacol. 175, 2825–2833 (2018). [PMC free article] [PubMed] [Google Scholar]

41. Azizbeigi R., Farzinpour Z., & Haghparast A. Role of orexin-1 receptor within the ventral tegmental area in mediating stress- and morphine priming-induced reinstatement of conditioned place preference in rats. Basic Clin Neurosci. 10, 373–382 (2019). [PMC free article] [PubMed] [Google Scholar]

42. Pantazis C.B., James M.H., O’Connor S., Shin N., & Aston-Jones G. Orexin-1 receptor signaling in ventral tegmental area mediates cue-driven demand for cocaine. Neuropsychopharmacology (2021). [PMC free article] [PubMed] [Google Scholar]

43. Maduna T. et al. Microglia express mu opioid Receptor: Insights from transcriptomics and fluorescent reporter mice. Frontiers in Psychiatry 9, (2019). [PMC free article] [PubMed] [Google Scholar]

44. Horvath R.J. & DeLeo J.A. Morphine enhances microglial migration through modulation of P2X4 receptor signaling. The Journal of Neuroscience 29, 998 (2009). [PMC free article] [PubMed] [Google Scholar]

45. Machelska H. & Celik M. Opioid receptors in immune and glial cells-implications for pain control. Frontiers in Immunology 11, (2020). [PMC free article] [PubMed] [Google Scholar]

46. Cady R.J., Denson J.E., Sullivan L.Q., & Durham P.L. Dual orexin receptor antagonist 12 inhibits expression of proteins in neurons and glia implicated in peripheral and central sensitization. Neuroscience 269, 79–92 (2014). [PubMed] [Google Scholar]

47. Li Y. & van den Pol A.N. Mu-opioid receptor-mediated depression of the hypothalamic hypocretin/orexin arousal system. Journal of Neuroscience 28, 2814–2819 (2008). [PMC free article] [PubMed] [Google Scholar]

48. Stoeber M. et al. A genetically encoded biosensor reveals location bias of opioid drug action. Neuron 98, 963–976 (2018). [PMC free article] [PubMed] [Google Scholar]

49. Peyron C. et al. A mutation in a case of early onset narcolepsy and a generalized absence of hypocretin peptides in human narcoleptic brains. Nat. Med.. 6, 991–997 (2000). [PubMed] [Google Scholar]

50. Thannickal T.C. et al. Reduced number of hypocretin neurons in human narcolepsy. Neuron. 27, 469–474 (2000). [PMC free article] [PubMed] [Google Scholar]

51. Thannickal T.C. et al. Human narcolepsy is linked to reduced number, size and synaptic bouton density in hypocretin-2 labeled neurons. Abstr Soc Neurosci 26, 2061 (2000). [Google Scholar]

52. James M.H., Mahler S.V., Moorman D.E., & Aston-Jones G. A decade of orexin/hypocretin and addiction: where are we now? Curr Top Behav Neurosci 33, 247–281 (2017). [PMC free article] [PubMed] [Google Scholar]

53. Nestler E.J. Cellular basis of memory for addiction. Dialogues Clin Neurosci 15, 431–443 (2013). [PMC free article] [PubMed] [Google Scholar]

54. Carrera-Canas C., de Andres I., Callejo M., & Garzon M. Plasticity of the hypocretinergic/orexinergic system after a chronic treatment with suvorexant in rats. Role of the hypocretinergic/orexinergic receptor 1 as an autoreceptor. Frontiers in Molecular Neuroscience 15, (2022). [PMC free article] [PubMed] [Google Scholar]

55. Kaushik M.K. et al. Induction of narcolepsy-like symptoms by orexin receptor antagonists in mice. Sleep DOI: 10.1093/sleep/zsab043, (2021). [PubMed] [CrossRef] [Google Scholar]

56. Sora I. et al. Mu opiate receptor gene dose effects on different morphine actions: evidence for differential in vivo mu receptor reserve. Neuropsychopharmacology 25, 41–54 (2001). [PubMed] [Google Scholar]

57. Paqueron X. et al. Is morphine-induced sedation synonymous with analgesia during intravenous morphine titration? British Journal of Anaesthesia 89, 697–701 (2002). [PubMed] [Google Scholar]

58. Ito D. et al. Microglia-specific localisation of a novel calcium binding protein, Iba1. Molecular Brain Research 57, 1–9 (1998). [PubMed] [Google Scholar]

